# MT5-MMP C-terminal mutations reduce C99 and Aβ accumulation: A foundation for peptide-based Alzheimer’s interventions

**DOI:** 10.1101/2025.05.22.655451

**Authors:** Pedro Belio-Mairal, Athina Kamitsou, Delphine Stephan, Nicolas Jullien, Diarra Thiane, Melissa Ramos, Laurence Louis, Florian Benoist, Baptiste Serrano, Marion David, Michel Khrestchatisky, Pascaline Lécorché, Emmanuel Nivet, Santiago Rivera

## Abstract

Our prior work established a pivotal role for membrane-type 5-matrix metalloproteinase (MT5-MMP) in Alzheimer’s disease (AD) pathogenesis, particularly through its C-terminal transmembrane (TM) and intracellular (IC) domains, which influence the fate of major toxic metabolites of toxic amyloid precursor protein (APP) metabolites, particularly C99 and Aβ. Hypothesizing that modifications in these domains could modulate C99 and Aβ levels, we engineered MT5-MMP variants with deletions or substitutions in the TM/IC domains or specific IC amino acid clusters. When co-transfected into human cell lines accumulating C99, certain IC domain mutations promoted C99 degradation and reduced Aβ levels, while other mutations had divergent effects. High content imaging further revealed that MT5-MMP IC domain modification altered C99 subcellular trafficking within the endomembrane system, impacting its processing. Proximity ligation assays confirmed the importance of the IC domain in MT5-MMP colocalization and potential interaction with C99. To translate these findings, we synthetized peptides mimicking the MT5-MMP IC domain, incorporating mutations that reduce C99 and/or Aβ levels. One peptide effectively lowered C99 levels in an *in vitro* AD model. Overall, this study highlights the importance of specific amino acids in the C-terminal domains of MT5-MMP for regulating C99 and Aβ metabolism. It also provides new insights for developing MT5-MMP-based therapeutic strategies against AD, exploiting the unique properties of specific mutations in MT5-MMP to prevent toxic accumulation of C99 and Aβ.

GRAPHICAL ABSTRACT

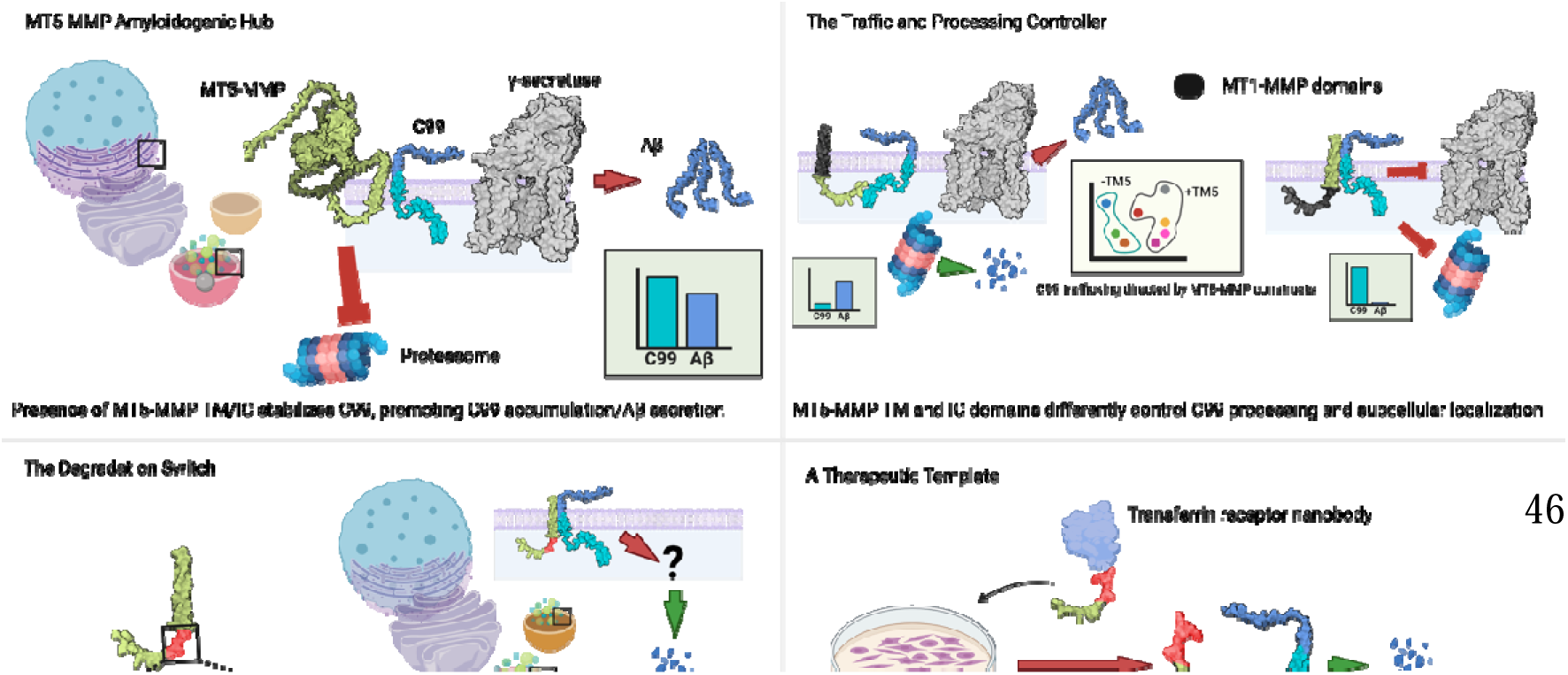

## INTRODUCTION

Matrix metalloproteinases (MMPs) form a multigenic family of more than 20 endopeptidases with a wide spectrum of biological functions^1^. Membrane-type 5-matrix metalloproteinase (MT5-MMP, also known as MMP-24 or η-secretase), is the only MMP primarily expressed in neural cells^2,3^ and plays a critical role in physiological and pathological processes in the nervous system, including: synaptic plasticity, neural stem cell differentiation, post-lesion synaptic reorganization, and axonal growth and regeneration^4,5^

The presence of MT5-MMP in amyloid plaques of Alzheimer’s disease (AD) patients suggests its involvement in AD pathogenesis^2^. Consistent with this, MT5-MMP cleaves amyloid precursor protein (APP) in HEK cells^6^ and murine primary neurons^7^. *In vivo* MT5-MMP deficiency reduces APP processing in the 5xFAD mouse model of AD^8,9^, while its overexpression increases C99 and Aβ accumulation in HEK cells carrying the Swedish familial AD mutation (HEKswe)^8^. APP processing is a central event in AD pathogenesis, whereby β-secretase cleavage generates the APP C-terminal fragment (CTF) C99, which γ-secretase further processes to release the beta-amyloid peptide (Aβ)^10^. Beyond its effects on APP metabolism, MT5-MMP deficiency reduces neuroinflammatory signaling and glial reactivity while preserving long-term potentiation (LTP) and cognitive functions in 5xFAD mice^8,9^.

Growing evidence highlights the importance of molecular events that occur upstream of Aβ pathology in AD. Accumulation of C99 precedes Aβ buildup in both the 3xTg and 5xFAD mouse models^11,12^, and C99 also accumulates in the brains of AD patients, where its levels correlate with neuronal vulnerability and cognitive impairment^13^. Importantly, C99 accumulation induces endolysosomal dysfunctions independently of Aβ in both mouse and human models, potentially explaining its early neurotoxic effects^11,14^. We previously reported that MT5-MMP regulates the metabolism of APP CTFs *via* both proteolytic-dependent and -independent mechanisms^4^. Notably, the MT5-MMP transmembrane (TM) and intracellular (IC) non-catalytic domains are sufficient to sustain high levels of C99 and Aβ in HEK cells overexpressing C99. Furthermore, a truncated MT5-MMP containing only these two domains coimmunoprecipitates with C99, suggesting an interaction between these proteins ^4^. Together, these findings support a key role for non-catalytic functions of MT5-MMP in the regulation of C99 metabolism and suggest that this enzyme may contribute to early pathogenic events occurring upstream of Aβ deposition.

In this study, we investigated the mechanisms by which non-catalytic TM and IC domains of MT5-MMP domains regulate C99 metabolism. Using complementary molecular, cellular, and structure-function approaches, we assessed the role of specific amino acid residues within these domains in controlling C99 and Aβ levels. Our findings reveal novel aspect of the functional interplay between MT5-MMP and C99 and provide mechanistic insights into how MT-MMP modulates C99 processing and trafficking, laying a foundation for therapeutic strategies that modulate C99 and Aβ by harnessing the non-catalytic properties of MT5-MMP.

## MATERIALS AND METHODS

### Plasmid constructs

All plasmids were constructed using a pcDNA3.1 backbone. The new plasmids used in this study (ΔEXT, TM1/IC5, TM5/IC1, and mutated TM/IC variants) (Fig. 1A, 2A) were generated by GeneArt (ThermoFisher Scientific). Other MT5-MMP variants (pcDNA, TM/IC, EXT/TM, EXT) and C99 or HA-C99 were produced, purified, and amplified as previously described^4^.

**Figure 1.**
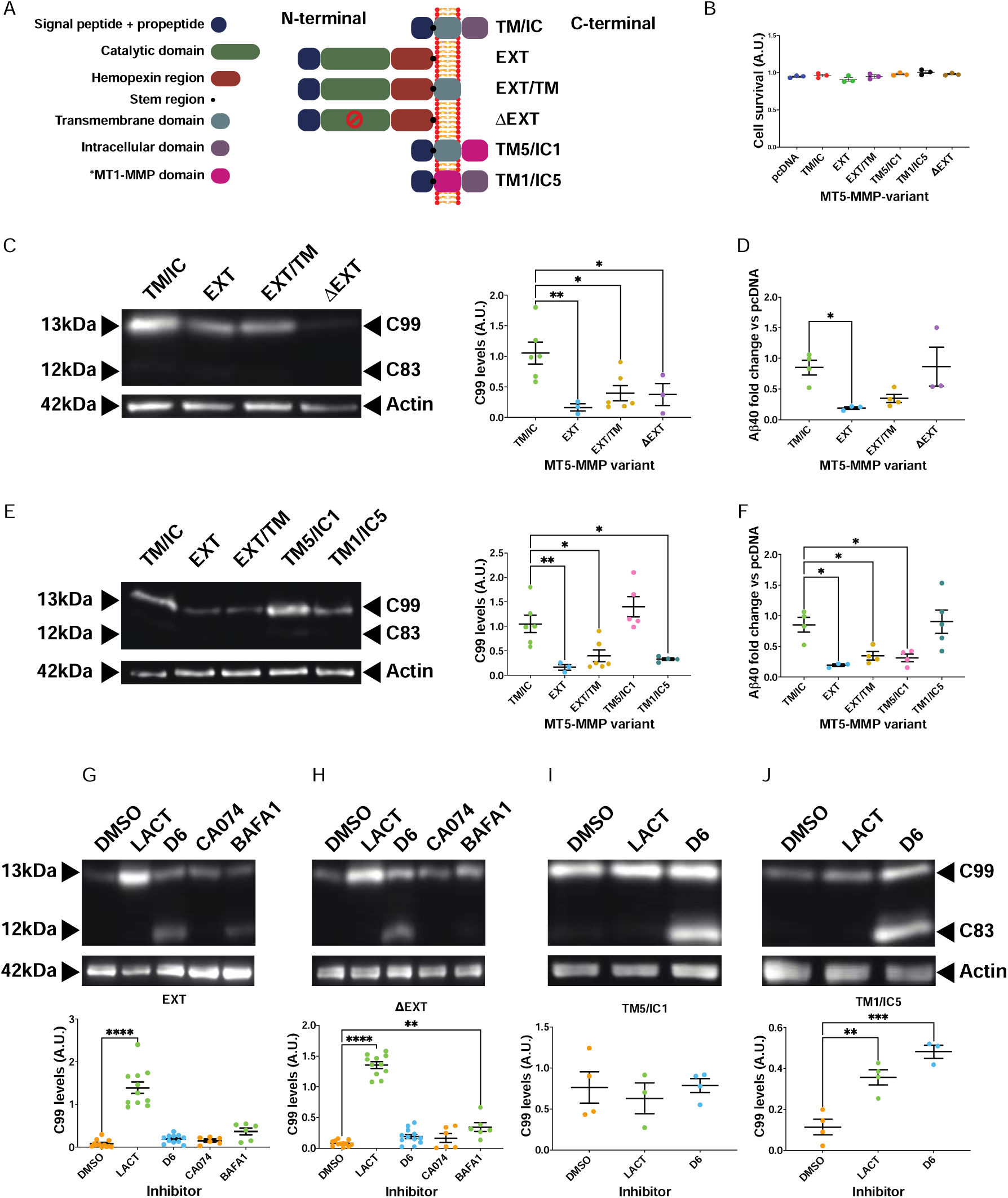
Whole-domain mutations modulate the accumulation of C99 and the production of Aβ in HEK cells. (A) Schematic representation of the MT5-MMP variants used. (B) Graph showing no changes in cell survival measured by the MTT assay 48 h after co-transfection of HEK cells with C99 and different MT5-MMP variants. Absorbance values are expressed in arbitrary units (A.U) and normalized to those of samples treated with transfection reagent alone. (C) Representative western blots showing C99 levels detected with an anti-APP-CTF antibody in HEK cells co-transfected with C99 and TM/IC, EXT, EXT/TM, or ΔEXT. The graph shows C99 levels normalized to actin as a protein loading control and the pcDNA condition. (D) Graphical representation of Aβ40 levels in the supernatants of HEK cells co-transfected with C99 and TM/IC, EXT, EXT/TM or ΔEXT. Values were normalized to the pcDNA condition for each experiment. (E) Representative western blots showing C99 levels detected with an anti-APP-CTF antibody in HEK cells co-transfected with C99 TM/IC, EXT, EXT/TM, TM5/IC1 or TM1/IC5. The graph represents C99 levels normalized to actin as a protein loading control and the pcDNA condition. (F) Graphical representation of Aβ40 levels in the supernatants of HEK cells co-transfected with C99 and with TM/IC, EXT, EXT/TM, TM5/IC1 or TM1/IC5. Values were normalized to the pcDNA condition for each experiment. (G-J) Representative western blots showing C99 levels detected with an anti-APP-CTF antibody in HEK cells co-transfected with C99 and (G) EXT, (H) ΔEXT, (I) TM5/IC1 or (J) TM1/IC5 variants. The corresponding actin-normalized quantifications are shown below each immunoblot. Cells were treated for 20 h with inhibitors of C99 processing pathways: proteasome (lactacystin; LACT 10 µM); γ-secretase ELND006 (D6 1 µM); cathepsin (B CA-074 Me; CA074 10 µM); and autophagolysosomal pathway (bafilomycin A1; BAFA 50 nM). DMSO was used as control vehicle. ANOVA followed by Dunnett’s post hoc test compared to control condition. Values are expressed as mean ± SEM of at least 3 independent experiments. P-values are shown as * p < 0.05, ** p < 0.01, *** p < 0.001, and **** p < 0.0001.

**Figure 2.**
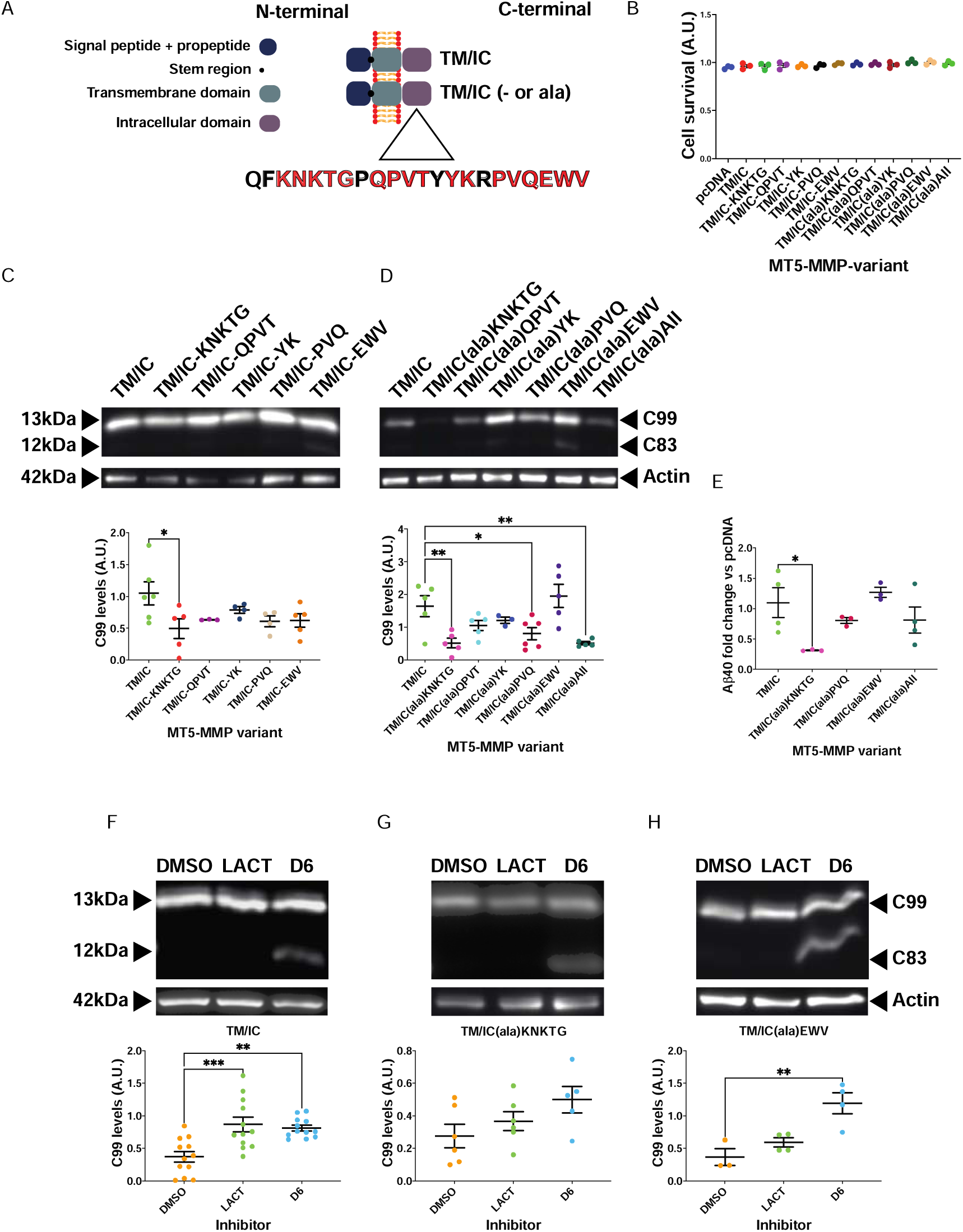
Mutation of specific MT5-MMP IC residues decreases C99 and Aβ levels in HEK cells. (A) Schematic representation of the MT5-MMP variants used. The amino acid groups in red (KNKTG, QPVT, YK, PVQ, and EWV) were either deleted or substituted by ala. (B) Graph showing no changes in cell survival measured by the MTT assay 48 h after co-transfection of HEK cells with C99 and different MT5-MMP variants. Absorbance values are expressed in arbitrary units (A.U) and normalized to those of samples treated with the transfection reagent alone. (C & D) Representative western blots showing C99 levels detected with an anti-APP-CTF antibody in HEK cells co-transfected with C99 and TM/IC or TM/IC with deletions (C) or ala substitutions (D) in KNKTG, QPVT, YK, PVQ or EWV. In addition, all amino acids were substituted with alanine (All) in D. The graphs below show the levels of C99 normalized to actin as a protein loading control and the pcDNA condition. (E) Graphical representation of Aβ40 levels in the supernatants of HEK cells co-transfected with C99 and TM/IC, TM/IC(ala)KNKTG, TM/IC(ala)PVQ, TM/IC(ala)EWV or TM/IC(ala)All. Values were normalized to the pcDNA condition for each experiment. (F-H) Representative western blots showing C99 levels detected with an anti-APP-CTF antibody in HEK cells co-transfected with C99 and (F) TM/IC, (G) TM/IC(ala)KNKTG or (H) TM/IC(ala)EWV and treated for 20 h with lactacystin (LACT 10 µM) or γ-secretase inhibitor (ELND006; D6 1µM). DMSO was used as a control vehicle. The graphs below represent C99 levels normalized to actin as a protein loading control. ANOVA followed by Dunnett’s post hoc test compared to control condition. Values are expressed as mean ± SEM of at least 3 independent experiments. P-values are shown as * p < 0.05, ** p < 0.01, and *** p < 0.001.

### Cell culture

HEK cells were maintained in DMEM Glutamax (61965026, ThermoFisher Scientific) supplemented with 10% fetal bovine serum (FBS) (A3160801, ThermoFisher Scientific) and 1% penicillin/streptomycin (15140122, ThermoFisher Scientific) (hereafter referred as HEK medium). For passaging, cells were washed with DPBS (14190094, ThermoFisher Scientific) and dissociated with 0.05% Trypsin-EDTA (25300054, ThermoFisher Scientific). After resuspension in HEK medium, cells were replated with fresh HEK medium containing Geneticin (10131027, ThermoFisher Scientific) at a 1:100 dilution.

hTERT-RPE1 cells were cultured similarly but in DMEM/F12 (31331028, ThermoFisher Scientific) supplemented with 10% FBS and 1% penicillin/streptomycin. HBSS (14025050, ThermoFisher Scientific) was used for washing and 0.25% Trypsin-EDTA (25200056, ThermoFisher Scientific) was used for cell dissociation.

### Transfections and treatments

Cells were transfected using JetPei^®^ Polyplus (101000053, Sartorius). For 96-well plates (6055300, Revvity; 353072, Corning), reverse transfection was used, with hTERT-RPE1cells plated at 1·10^4^ cells/well. For 6-well plates (353046, Corning), HEK cells were plated at 1.5·10^6^ cells/well and the classical forward transfection was performed. All transfections followed the manufacturer’s recommended volumes for transfection reagents and DNA. Synthetic peptides were cotransfected with DNA plasmids.

Proteinase and ATPase inhibitors (Table 1) were diluted in DMSO (D2650, Sigma-Aldrich) and added 15–17 h post-transfection. After 20 h of treatment, cells were lysed and collected for analysis.

**Table 1.** List of the inhibitors with concentration of the stock and final solutions, and manufacturer’s references.

| **Inhibitors** |  |  |  |  |
| --- | --- | --- | --- | --- |
| **Inhibitor** | **[Stock](μM)** | **Dilution** | **Reference** | **Company** |
| **LACT** | **50000** | **1:5000** | **BML-PI104-1000** | **Enzo Life Sciences** |
| **D6** | **5000** | **1:5000** | **SML2122** | **Sigma-Aldrich** |
| **BAFA1** | **250** | **1:5000** | **B1793** | **Sigma-Aldrich** |
| **CA074** | **50000** | **1:5000** | **205531** | **Sigma-Aldrich** |

### Cell viability assay (MTT)

Cell viability was assessed 48 h post-transfection or peptide treatment using the 3-(4,5-dimethylthiazol-2yl)-2,5-diphenyl tetrazolium bromide (MTT) assay (M5655 Sigma-Aldrich). MTT was diluted in HEK medium and added to wells to a final concentration of 0.5 mg/mL for 3 h at 37°C and 5% CO_2_. After incubation, the culture medium was replaced with 100 µL of dimethyl sulfoxide (DMSO) (D2650, Sigma-Aldrich) to dissolve the formazan crystals formed in the mitochondria of living cells. The absorbance was measured in a multiwell plate reader and viability was calculated as the percentage of live cells = (transfected cell OD_550_/control cell OD_550_) x 100, after normalizing the values by column and row.

### Protein quantification

Protein concentrations were determined using a BCA assay kit (23225, ThermoFisher Scientific) was used following the manufacturer’s instructions for the multiwell plate format. Standard curves were generated using a BSA standard kit (23208, ThermoFisher Scientific). Absorbance from two technical replicates was measured using a multiwell plate reader and concentrations were extrapolated using the 4-parameter fit of MARS data analysis software version 2.10R3 (BMG Labtech). All measurements were performed in duplicate with the same software.

### Western Blot

Samples with 20 µg of protein were mixed with 2x Tris-Glycine (LC2676, ThermoFisher Scientific) or Tricine (LC1676, ThermoFisher Scientific) sample buffer and heated at 95°C for 5 min or 85°C for 20 min for proteins <20 kDa.

Proteins were separated using a Mini gel tank system (A25977, ThermoFisher Scientific) and a Power Blotter Semi-dry Transfer system (PB0013, ThermoFisher Scientific) and Novex 4-20% Tris-Glycine (10- or 15-well, XP04200BOX/XP04205BOX) or Novex 16% Tricine gels (10- or 15-well, EC6695BOX/EC66955BOX), all from ThermoFisher, and run following the manufacturer’s instructions. Gels were transferred into 0.45 µm nitrocellulose membranes (10600002, Amersham) using a Power Blotter Semi-dry Transfer system (PB0013) in Towbin buffer.

For immunoblotting, membranes were stained with Ponceau (P7170, Sigma-Aldrich) for 30 sec and imaged using the Preview mode on a NineAllience 9.7 17.02 (UVITec). After blocking for 1 h in 1x PBS (ET330-A, Euromedex) with 5%_m/v_ powder milk and 0.2%_v/v_ Tween20 (P1379, Sigma-Aldrich), membranes were incubated overnight at 4°C under agitation with primary antibodies diluted (1:1000; anti-actin at 1:5000) in blocking buffer. After washing, membranes were incubated with HRP-conjugated secondary antibodies (1:1000) in blocking buffer for 1 h at RT. Immunoreactive bands were visualized using ECL (RPN2106, Amersham) or ECL Prime (RPN2236, Amersham), using the chemiluminescence mode of the UVITec station, and semi-quantified using Fiji’s “Plot lanes” tool (Gels module)^15^. Actin served as the loading and normalizing control.

### Immunocytochemistry

Cells were cultured in 96-well plates (6055300, Revvity) for high content screening microscopy (HCS). For sample fixation, PEM buffer was prepared to obtain a final solution of PIPES 80 mM (P1851, Sigma-Aldrich), MgCl_2_ 2mM (M9272, Honeywell), and EGTA 5mM (E4378, Sigma-Aldrich) at pH 6.8 in ddH_2_O. Then, paraformaldehyde (15714, Electron Microscopy Services) was diluted at 4%_v/v_ in PEM with 4%_m/v_ sucrose (S0389, Sigma-Aldrich) (fixation solution).

Culture media was removed from 96-well plates and then washed with 1x DPBS. After fixing with fixation solution for 10 min at RT, cells were washed 4 times with DPBS, then blocked and permeabilized for 1 hour at RT with 3%_m/v_ bovine serum albumin (BSA) (A7906, Sigma-Aldrich), 0.1% saponin (84510, Sigma-Aldrich), and 10% normal goat (005-000-121, Jackson ImmunoResearch) or donkey (017-000-121, Jackson ImmunoResearch) serum in 1x DPBS (block/per buffer). Primary antibodies were added to cells and incubated overnight at 4°C at the corresponding concentrations (Table 2). The next day, cells were washed with 1x DPBS and the Alexa Fluor secondary antibodies diluted in blocking/permeabilizing buffer (Table 2) incubated for 1 hour at RT. After washing in 1x DPBS cells were incubated for 10 min at RT with 10 µg/mL Hoechst33342 (H3570, ThermoFisher Scientific) diluted in 1x DPBS. After washing twice with 1x DPBS, plates were sealed with parafilm and stored in 1xDPBS. Plates were imaged within 24 h.

**Table 2.** List of antibodies with target antigens, host species, working dilutions, references, and the application.

| **Antibodies** |  |  |  |  |  |
| --- | --- | --- | --- | --- | --- |
| **Target** | **Host** | **Dilution** | **Reference** | **Company** | **Application** |
| **Primaries** |  |  |  |  |  |
| **APP-CTF** | **Rabbit** | **1:1000** | **A8717** | **Sigma-Aldrich** | **Western blot** |
| **Actin** | **Mouse** | **1:5000** | **A1978** | **Sigma-Aldrich** | **Western blot** |
| **Nogo-B** | **Sheep** | **1:25** | **AF6034** | **Biotechne** | **Immunofluorescence** |
| **GM130** | **Mouse** | **1:200** | **610823** | **BD Biosciences** | **Immunofluorescence** |
| **EEA1** | **Mouse** | **1:100** | **sc-365652** | **Santa Cruz Biotechnology** | **Immunofluorescence** |
| **LAMP1** | **Rabbit** | **1:300** | **ab24170** | **Abcam** | **Immunofluorescence** |
| **HA** | **Rat** | **1:100**<br>**1:100** | **11867423001** | **Roche** | **Immunofluorescence**<br>**PLA** |
| **Flag** | **Rabbit** | **1:200** | **14793** | **Cell Signaling Technology** | **PLA** |
| **Secondaries** |  |  |  |  |  |
| **Rabbit** | **Goat** | **1:800** | **111-005-003** | **Jackson ImmunoResearch** | **PLA** |
| **Rat** | **Goat** | **1:800** | **112-005-003** | **Jackson ImmunoResearch** | **PLA** |
| **Mouse-647** | **Donkey** | **1:800** | **A32787** | **ThermoFisher Scientific** | **Immunofluorescence** |
| **Rabbit-647** | **Donkey** | **1:800** | **A32795** | **ThermoFisher Scientific** | **Immunofluorescence** |
| **Sheep-647** | **Donkey** | **1:800** | **A21448** | **ThermoFisher Scientific** | **Immunofluorescence** |
| **Rat-488** | **Donkey** | **1:800** | **A48269** | **ThermoFisher Scientific** | **Immunofluorescence** |
| **Rabbit-488** | **Donkey** | **1:800** | **A-21206** | **ThermoFisher Scientific** | **Immunofluorescence** |
| **Mouse-594** | **Donkey** | **1:800** | **A-21203** | **ThermoFisher Scientific** | **Immunofluorescence** |

PLA stands for proximity ligation assay.

### Microscopy and data analysis

Images were acquired using the Operetta^©^ CLS High-Content Analysis System (Revvity). Image segmentation and feature extraction were performed with Harmony software (Revvity). The workflow for image acquisition, filtering, segmentation, and quantification is available at: github.com/RiveraLabINP7051/BelioMairal_2024. The resulting quantitative data were exported and subjected to downstream analysis in Python.

Python procedures to clean, downsize, normalize, and visualize quantitative data from the automated microscope were run in Jupyter Notebooks (available at: github.com/RiveraLabINP7051/BelioMairal_2024). In brief, overlap values for individual cells were multiplied by the HA fluorescence area to generate a new variable representing the total area of overlap of HA with each organelle. The overlap areas were normalized using Python’s sklearn. Individual cells were then clustered, and the dataset was downsized by excluding the clusters of the most extreme values. The mean area of overlap was calculated for each MT5-MMP variant and normalized to pcDNA. Finally, outliers were detected with the Tukey method and the values were used to visualize the results and perform Bayesian statistics.

After averaging the values by plasmid and organelle, the PCA function from sklearn was used with two principal components (n components=2). For cluster analysis, the function clustermap from the seaborn package was used.

A complete list of Python and package versions is provided at: github.com/RiveraLabINP7051/BelioMairal_2024.

### ELISA

Secreted Aβ40 in cell supernatants were measured using an ELISA kit (KHB3481, ThermoFisher Scientific). Supernatants were collected, stored at −80°C until analysis, and diluted to match the detection range before analysis, following the manufacturer’s protocol. All values were normalized to the the pcDNA condition.

### Proximity ligation assay

Colocalization between C99 and MT5-MMP constructs was assessed by proximity ligation assay (PLA) using the DuoLink^©^ In Situ system (Sigma-Aldrich). Procedures were adapted for 96-well plates. Unconjugated toat anti-rat (112-005-003, Jackson ImmunoResearch) and goat anti-rabbit (111-005-003, Jackson ImmunoResearch) antibodies were conjugated with DuoLink^©^ PLA orange probes (DUO96030-1KT, Sigma-Aldrich) following the manufacturer’s instructions and stored at 4°C. Cells were stained using the immunofluorescence protocol as described above. Secondary antibody binding and detection were performed using the DuoLink^©^ In situ orange detection kit (DUO92007-100RXN, Sigma-Aldrich). Cell nuclei were stained with Hoechst 33342 as described above.

### Peptide synthesis

Fmoc–amino acids were obtained from Iris Biotech (Marktredwitz, Germany). All other amino acids and reagents were purchased from Sigma-Aldrich (Saint-Quentin-Fallavier, France) or Analytical Lab (Castelnau-le-Lez, France). Peptide assembly was performed using a Liberty (CEM®) microwave synthesizer by solid phase peptide synthesis (SPPS) based on the Fmoc strategy. Fmoc–Gly-Wang resin (loading 0.66 mmol/g) were purchased from Iris Biotech and were used as solid support.

#### Purification and analysis of peptide-based products

Monitoring of reactions and quality controls of the peptide-based intermediates were carried out by liquid chromatography/mass spectrometry (LC/MS) using a Thermo Fisher Scientific UltiMate®3000 system (Waltham, MA, USA) equipped with an ion trap (LCQ Fleet) and an electrospray ionization source (positive ion mode). The LC flow rate was set to 2 mL/min using H2O 0.1%TFA (buffer A) and MeCN 0.1%TFA (buffer B) as eluents. The gradient elution was 10%–90% B in 10 min (quality control) equipped with a C18 Kinetex™ (2.6 μm, 50 mm × 4.6 mm). The heated electrospray ionization source had a capillary temperature of 350 °C. Crude peptides were purified using reverse-phase (RP)-High Pressure Liquid Chromatography (HPLC) on a Thermo Fisher Scientific UltiMate®3000 system equipped with a C18 Luna™ (5 μm, 100 mm × 21.2 mm). The detection was performed at 214 nm. The elution system comprised H2O 0.1%TFA (buffer A) and MeCN 0.1%TFA (buffer B). Flow rate was 20 mL/min.

#### Preparation of p17 and p20

Peptide p17: H-QFAAAAAPQPVTYYKRPVQEWV-COOH and peptide p20: H-QFKNKTGPQPVTYYKRAAAEWV-COOH were synthesized using 0.1 mmol SPPS Fmoc–Gly-Wang resin under microwave activation. Resin was swollen in DMF for 10 min. Deprotection of the fmoc-protecting group was performed using a solution of 20% piperidine in DMF for 200 s under microwave activation at 75 °C. Stepwise assembly of the Fmoc–protected-amino-acids were performed under microwave activation using standard Fmoc peptide chemistry. Double coupling times of 240 s were used with a solution of Fmoc–amino acid (1 eq), Oxyma (10 eq excess; 1 M) in DMF, and DIC (5 eq excess; 1 M) in DMF under microwave activation at 90 °C. Double coupling was used for R amino acids to improve yield. Finally, the peptidyl-resin was washed 3 times with DCM and then treated with TFA/TIS/H2O (95:2.5:2.5) containing DTT (10 mg/1mL) at RT for 4 h. The cleavage solution was recovered, concentrated under N2 flow, precipitated with cold diethyl ether and washed 2 times with cold diethyl ether. The crude products were dissolved in an H2O + 0.1%TFA /MeCN + 0.1% TFA (1:1) mixture and lyophilized. The crude peptide was purified using preparative RP-HPLC as described previously to obtain p17 and p20 as white powders (p17: 60 mg, 24%; p20: 52 mg, 20%).

### Statistics

Frequentist statistics were performed using GraphPad Prism 10.2.3 for MacOSX (GraphPad Software). Unless otherwise stated, ANOVA with Dunnett’s post hoc multiple comparison test, was used to compare all conditions to the control group. Graphs were generated by Prism, with p-values indicated as: * p value < 0.05, ** p value < 0.01, *** p value < 0.001, and **** p value < 0.0001.

For Bayesian statistics, JASP 0.18.3 (JASP team) was used to perform Bayesian ANOVA with post hoc tests. Prior odds were the software default values, and the Bayes factor was calculated comparing the alternative (1) against the null (0) model (BF10). The software directly provided the logarithmic BF10 (LogBF10).

All statistical analyses were performed with at least three independent experiments.

## RESULTS

### Modulation of C99 and Aβ levels by different MT5-MMP variants

We previously showed that deletion of the intracytoplasmic (IC) domain of MT5-MMP alone (EXT/TM) or together with the transmembrane (TM) domain (EXT), significantly reduced C99 levels in HEK cells coexpressing C99. In contrast, the TM/IC variant, lacking the extracellular domains, reproduced the pro-amyloidogenic effects of MT5-MMP in C99-overexpressing cells^4^. To further elucidate the role of specific domains or sequences in regulating C99 and Aβ levels, we generated plasmids encoding mutant forms of MT5-MMP (Fig. 1A) and cotransfected them with C99 in HEK cells. None of these cotransfections affected cell viability, as measured by the MTT assay (Fig. 1B).

To determine whether the proteolytic activity of the extracellular (EXT) domain could account for its anti-amyloidogenic effects, we generated an inactive EXT variant (ΔEXT) by replacing the Glu283 residue of the catalytic site with alanine. Coexpression of EXT, EXT/TM, and ΔEXT in HEK cells reduced C99 levels by 83%, 71%, and 74%, respectively, compared to the TM/IC control (Fig. 1C). This indicates that the catalytic activity of EXT and EXT/TM is not required for C99 decrease in the absence of TM and IC domains. Notably, EXT expression significantly decreased Aβ levels in cell supernatant (ELISA, Fig. 1D), whereas ΔEXT maintained Aβ levels close to control (Fig. 1D). This suggests that inactivation of the catalytic domain may redirect C99 toward Aβ production, unlike the active EXT variant.

To determine whether changes in C99 were primarily driven by the TM or IC domain, we generated chimeric constructs by swapping domains between MT5-MMP and its close homologue, MT1-MMP. These substitutions were based on functional differences between the two proteases: suppression of MT1-MMP IC domain increases C99 levels, whereas suppression of the MT5-MMP IC domain decreases them^4^. We created two chimeras: TM5/IC1 (MT5-MMP TM + MT1-MMP IC) and TM1/IC5 (MT1-MMP TM + MT5-MMP IC) (see Fig. 1A). TM5/IC1 maintained C99 levels but reduced Aβ secretion by 67% (Fig. 1E, F). In contrast TM1/IC5 significantly decreased C99 levels by 69% while maintaining Aβ levels comparable to TM/IC control (Fig. 1E, F). These contrasting results highlight the distinct influence of MT5-MMP TM and IC domains on C99 processing and Aβ production.

### MT5-MMP non-catalytic variants control C99 processing

We next investigated the fate of C99 by modulating enzymatic systems potentially involved in its degradation, including the proteasome (inhibited by lactacystin, LACT), γ-secretase (inhibited by ELND006, D6), the autophagolysosome system (inhibited by Bafilomycin A1, BAFA1), and cathepsin B (inhibited by CA-074-Me, CA074), which has no effect on canonical APP secretases^16^.

In EXT-expressing cells, only LACT treatment caused a significant recovery of C99 levels, whereas all other inhibitors had no effect (Fig. 1G), indicating a prominent role of the proteasome in C99 degradation upon coexpression with EXT. Similarly, in the ΔEXT group, LACT also induced a significant recovery of C99, whereas D6 and CA074 had no effect (Fig. 1H). Considering that coexpression of ΔEXT and C99 maintains Aβ levels, the failure of D6 to recover C99 suggests that EXT catalytic activity hinders Aβ generation. BAFA1 treatment induced a significant 331% recovery of C99 levels in ΔEXT-cotransfected cells, suggesting that the inactivation of the MT5-MMP catalytic domain partially redirects C99 to lysosomal compartments. These data indicate that the proteasome is the primary processing machinery of C99 in the presence of EXT or ΔEXT, with an alternative degradation pathway involving EXT catalytic activity in lysosomal compartments. CA074 failed to recover C99 levels in EXT- or ΔEXT-expressing cells (Fig. 1G, H), suggesting that the inhibition of this lysosomal enzyme is insufficient to restore C99 levels and block Aβ production.

The prominent role of the proteasome in C99 processing in the presence of MT5-MMP TM and IC prompted us to compare its modulation with γ-secretase, the main processing pathway converting C99 in Aβ. Analyses of these two pathways in the presence of each chimera yielded distinct results: LACT and D6 had no effect on C99 levels in TM5/IC1-expressing cells (Fig. 1I). In contrast, in cells expressing the TM1/IC5 variant, LACT and D6 increased C99 levels by 212% and 323%, respectively (Fig. 1J), suggesting that both the proteasome and γ-secretase mediate the reduction in C99 levels mediated by this chimera.

### A specific cluster of amino acid deletions in the IC domain induces a reduction in C99 and Aβ levels

We next investigated whether the deletion of small clusters of residues in the MT5-MMP IC domain could replicate the reduction in C99 levels caused by full IC domain deletion. As mentioned above, the selected deletions—KNKTG, QPVT, YK, PVQ, and EWV (Fig. 2A)— were based on sequence differences between the IC domains of MT5-MMP and MT1-MMP^4^, which may underlie their opposing effects on C99 levels upon IC domain deletion. None of the mutant variants affected cell survival in HEK cells coexpressing C99 (Fig. 2B). While the deletions generally reduced C99 levels by 40% ± 10%, only the KNKTG deletion reached statistical significance, reducing C99 levels by 53% (Fig. 2C).

### Sequence substitutions in the IC domain modulate C99 and Aβ levels

The limited impact of targeted deletions on C99 levels may imply potential conformational changes in the IC domain that could interfere with the physiological effects of MT5-MMP variants. To address this, we introduced alanine (Ala) substitutions in the same clusters and included a construct where all selected amino acids were simultaneously replaced with ala (All). None of these substitutions affected cell viability (Fig. 2B). When coexpressed with C99 in HEK cells, only (Ala)KNKTG or (Ala)PVQ variants significantly reduced C99 levels by 69% and 51%, respectively, compared to the TM/IC control, as shown by Western blot using the anti-CTF antibody (Fig. 2D). Likewise, the All variant caused a significant 69% reduction in C99 levels (Fig. 2D). In contrast, Ala substitutions in the PDZ-binding EWV motif preserved C99 levels equivalent to those of TM/IC (Fig. 2D).

Among all the MT5-MMP with the potential to regulate C99 levels, only the KNKTG substitution significantly decreased Aβ levels by 72% (Fig. 2E), highlighting its prominent role in preventing C99 and Aβ accumulation. Conversely, the (Ala)EWV maintained high levels of both C99 and Aβ, comparable to the TM/IC control.

To investigate whether the reduction in C99 levels after (Ala)KNKTG expression resulted from altered proteolytic degradation, we treated HEK cells with LACT and D6 after cotransfection with C99 and either TM/IC, TM/IC(ala)KNKTG, or TM/IC(ala)EWV. LACT and D6 significantly restored C99 levels when cotransfected with TM/IC (Fig. 2F). However, in the TM/IC(ala)KNKTG group, C99 levels remained unchanged upon treatment with LACT or D6 (Fig. 2G), suggesting an alternative degradation pathway independent of the proteasome or γ-secretase. In HEK cells cotransfected with TM/IC(ala)EWV, only D6 increased C99 levels (Fig. 2I), indicating that Ala substitution in the EWV motif favors γ-secretase-mediated processing of C99 over proteasomal degradation. These findings suggest that the significant changes in C99 fate associated with MT5-MMP variants may reflect their translocation to specific subcellular compartments, facilitating C99 processing by distinct proteolytic systems.

### MT5-MMP variants alter the pattern of subcellular localization of C99

To investigate how MT5-MMP variants influence the subcellular distribution of C99, and its metabolic fate, we co-transfected human retinal pigment epithelial (RPE1) cells with HA-N-terminally tagged C99 (HA-C99) and MT5-MMP variants known to differentially modulate C99 processing (Fig. 3A). RPE1 are highly adherent and optimally sized for analyzing subcellular protein distribution using high-throughput confocal microscopy. Additionally, these cells do not express detectable levels of MT5-MMP (Fig. 3B) and, like neural cells, are of embryonic neuroectodermal origin^17^. We used anti-HA antibodies to detect HA-C99 and combined them with antibodies targeting endomembrane organelles (early endosomes, Golgi apparatus, endoplasmic reticulum (ER), and lysosomes) to study HA-C99 subcellular localization via immunofluorescence. Images were acquired and analyzed using quantitative unbiased high-content screening (HCS), with pcDNA serving as a reference to calculate HA distribution changes as a ratio of MT5-MMP variants to pcDNA.

**Fig 3.**
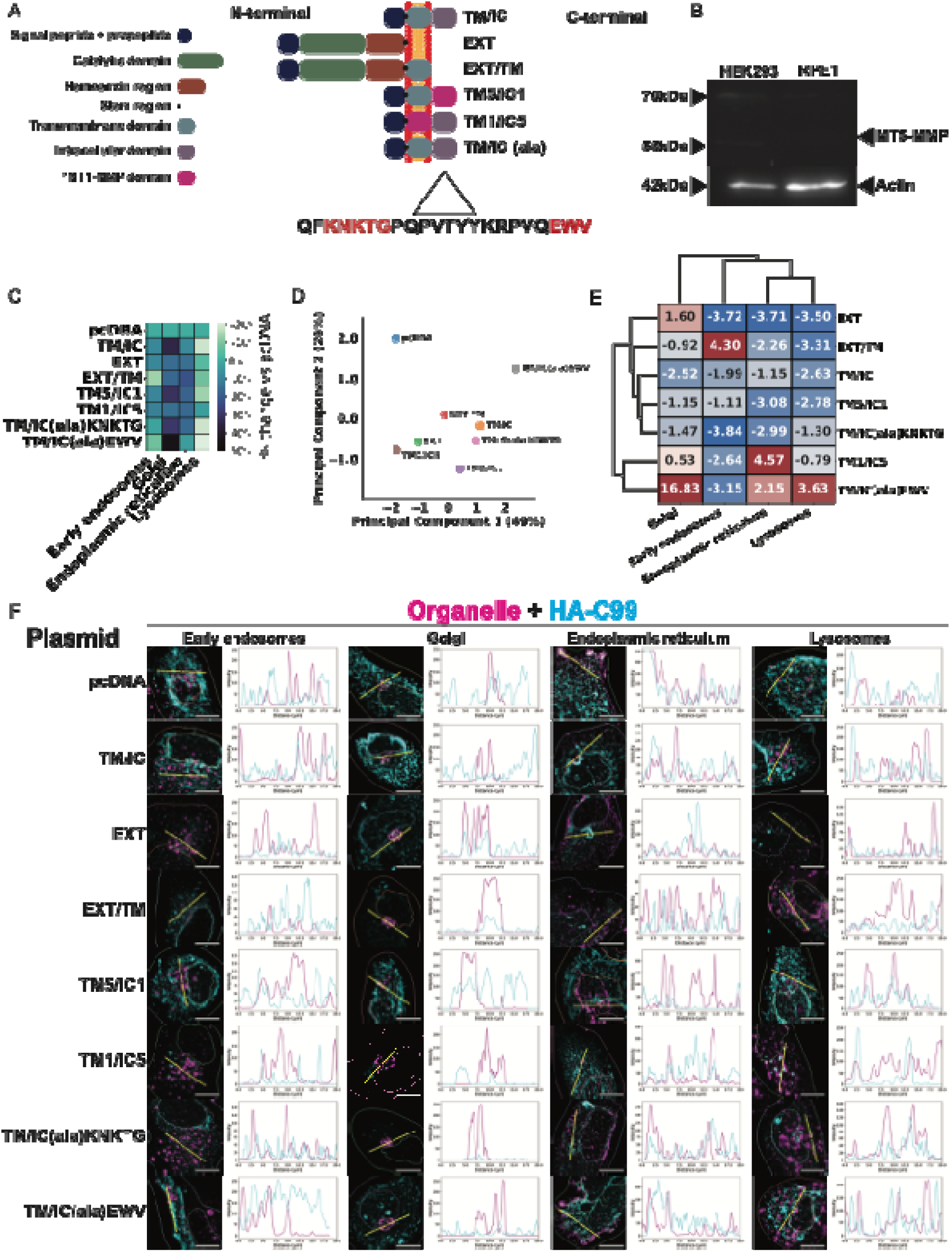
Effect of specific MT5-MMP IC domain mutations on the subcellular distribution of C99 in RPE1 cells. (A) Schematic representation of the MT5-MMP used. The amino acid groups in red (KNKTG and EWV) were substituted by ala. (B) Western blotting showing the lack of MT5-MMP expression in lysates of both HEK and RPE1 cells. (C) Heatmap representing the quantification of the area of HA^+^ signal in RPE1 cells overlapping with each organelle as a function of the MT5-MMP variant co-transfected with HA-C99 and relative to pcDNA in percent change. (D) Principal component analysis (PCA) of the quantification heatmap in C, constructed with two components representing 75% of the total variability of the data. Each MT5-MMP variant clustered with variants that had similar profiles of change relative to pcDNA. (E) Cluster map representing changes in (D) based on Bayesian ANOVA. The values are the summed LogBF10 values of multiple comparisons between MT5-MMP variants in each organelle, representing the statistical strength of variability in HA overlap with organelles for each MT5-MMP variant. The higher the values, the stronger the statistical evidence that a given MT5-MMP variant induces changes in HA overlap relative to all other variants in the same organelle. The lower the values, the weaker the statistical evidence for such changes. Higher values are shown in light to dark red, and lower values are represented in light to dark blue. In this cluster map, TM/IC(ala)EWV particularly affects HA overlap in the Golgi, ER, and lysosomes relative to all other MT5-MMP variants; and TM1/IC5 specifically affects HA signal overlap with the ER. However, compared with the other variants, EXT did not alter the HA overlap with lysosomes compared to the other variants. (F) Representative HCS fluorescence micrographs of HA signal overlapping with organelle immunolabeling in RPE1 co-transfected with HA-C99 and MT5-MMP variants. The organelles are shown in magenta, and the HA signal is shown in cyan. The multiline plots show the distribution of fluorescence intensity along the 20 µm yellow line in the image, representing the colocalization of fluorescence along the line. Scale bar: 10 µm.

The distribution of the HA signal varied under the influence of different MT5-MMP variants (Fig. 3C). Notably, all MT5-MMP variants caused an overall reduction in HA signal within the Golgi apparatus (27%±12) and the ER (14%±6) relative to the pcDNA control. In contrast, we observed inconsistent changes in HA signal among MT5-MMP variants in early endosomes (10%±10), accompanied by a slight increase (11%±9) in the lysosomal HA signal (Fig. 3C). However, absolute quantification is insufficient to assess the complexity of HA signal distribution across organelles under the influence of different MT5-MMP variants. To better visualize these effects, we conducted a principal component analysis (PCA) using two principal components, which collectively account for 75% of HA signal variability (Fig. 3D). Both pcDNA and TM/IC(ala)EWV were isolated from the main cluster of MT5-MMP variants, indicating that coexpression of either construct leads to unique HA-C99 distribution patterns. In addition, three distinct groups emerged within the main cluster: variants with MT5-MMP TM (EXT/TM, TM/IC, TM/IC(ala)KNKTG), variants without MT5-MMP TM (EXT, TM1/IC5), and the TM5/IC1 chimera, positioned between these groups. These results suggest that: i) HA-C99 distribution across organelles varies depending on the coexpression of MT5-MMP variants, and ii) The presence of MT5-MMP TM determines the subcellular localization of HA-C99.

To assess the effect of MT5-MMP constructs on HA-C99 distribution in each organelle, we performed cluster analysis using the results of Bayesian ANOVA followed by multiple comparison tests. The natural logarithm of Bayes Factor 10 (LogBF10) quantifies the likelihood of differences in means between groups, providing a measure of the posterior probability of the alternative over the null hypothesis^18^. For example, a LogBF10 value of −1.1 infers that the null hypothesis is three times more likely, a LogBF10 of 0 indicates equal probability for both hypotheses, and a LogBF10 of 1 implies the alternative hypothesis is about three times more probable. Unlike conventional frequentist methods that rely on p-values, the Bayesian analysis provides a more quantitative criterion for evaluating group differences. In our results, LogBF10 provides a quantitative framework for interpreting whether variations in HA signal quantification are sufficient to support a difference in means or if the evidence remains inconclusive (e.g., due to high variability or small sample size).

We calculated the sum of LogBF10 values for each comparison of the effect of individual MT5-MMP variants on HA-C99 signal in each organelle to create a cluster map (Fig. 3E, F). This map that visually represents the strength of evidence for variability in different constructs across multiple organelles. TM/IC(ala)EWV differed from other constructs in HA signal distribution within the Golgi (16.83), ER (2.15), and lysosomes (3.63). However, other MT5-MMP variants altered HA-C99 distribution in specific cellular compartment: TM1/IC5 in the ER (4.57), EXT/TM in the endosomes (4.30), and EXT in the Golgi (1.60) (Fig. 3E, F).

### MT5-MMP variants heterogeneously determine HA-C99 subcellular localization

Bayesian analysis indicated that all MT5-MMP constructs altered the subcellular distribution of HA-C99 to some extent compared to the pcDNA control, with certain variants standing out. To explore these differences further, we analyzed LogBF10 values for individual comparisons in the post-hoc analysis (Fig. 4A-E).

**Figure 4.**
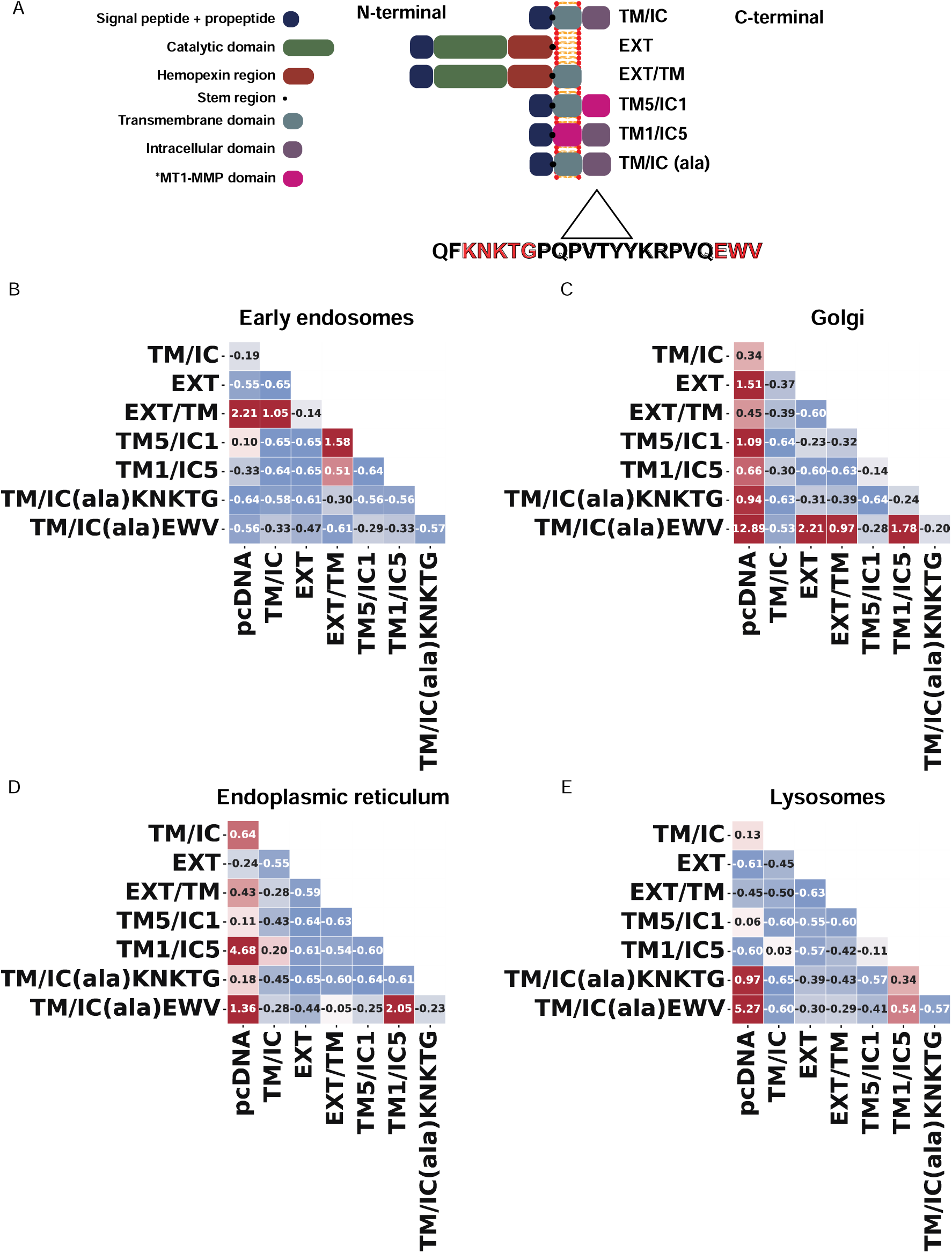
LogBF10 values of individual differences in the subcellular localization of HA-C99 in RPE1 cells transfected with different MT5-MMP variants. (A) Schematic representation of the MT5-MMP used. Amino acid groups in red (KNKTG and EWV) were replaced by ala. (B-E) Diagonal correlation matrices represent LogBF10 values for each comparison in a Bayesian ANOVA, with post-hoc multiple comparisons of the percentage of change in HA overlap induced by different MT5-MMP variants relative to pcDNA in early endosomes (B), Golgi (C), endoplasmic reticulum (D) and lysosomes (E). Higher levels are shown in light to dark red, and lower values are shown in light to dark blue. For instance, in (C) Golgi, the LogBF10 value from comparing pcDNA to TM/IC(ala)EWV indicates a significant reduction in HA signal in the Golgi. Values close to 0 indicate that there is insufficient evidence to determine whether there is a difference. Negative values represent strong evidence supporting no difference in the comparison of means.

Co-staining of EEA1 and HA (HA-C99) showed subtle changes in HA-C99 levels in early endosomes. Only EXT/TM increased (+13%) HA-C99 translocation into early endosomes compared to pcDNA (LogBF10 = 2.21), TM/IC (LogBF10 = 1.05), and TM5/IC1 (LogBF10 = 1.58) (Fig. 3F, 4D). This suggests that MT5-MMP EXT and/or TM, but not MT5-MMP IC, favors the presence of HA-C99 in EEA1^+^ vesicles.

We then examined the Golgi apparatus, a key organelle for protein packaging and compartmentalization. All MT5-MMP variants induced a significant reduction (between 15% and 45%) in HA-C99 compared with pcDNA (Fig. 3F, 4C). Among them, TM/IC(ala)EWV showed the strongest effect, with a significant 45% reduction and a LogBF10 of 12.89. In addition, HA-C99 reduction in the Golgi was more pronounced in TM/IC(ala)EWV (−45%) than in EXT (−16%) (LogBF10 = 2.21), EXT/TM (−17%) (LogBF10 = 0.97), and TM1/IC5 (−15%) (LogBF10 = 1.78). This suggests that the presence of MT5-MMP TM associated with an IC structure facilitates the translocation of HA-C99 outside the Golgi apparatus.

We next investigated whether the effects of MT5-MMP variants might occur upstream of the Golgi, in the ER, the primary site of protein synthesis. All MT5-MMP variants differentially reduced HA-C99 levels in the ER compared to pcDNA (Fig. 3F, 4B). In particular, TM1/IC5 reduced HA-C99 levels in the ER by 17% compared to pcDNA (LogBF10 = 4.68), a greater reduction than TM/IC(ala)EWV (−4%) (LogBF10 = 2.05). These differences in HA signal in the ER suggest that the MT5-MMP variants induce the exit of HA-C99 to other cellular compartments, with some variants being more effective than others.

Based on our previous results, the levels of co-localization between early endosomes and HA-C99 indicate that EXT/TM preferentially transports HA-C99 from the ER/Golgi to early endosomes, whereas other MT5-MMP variants may translocate HA-C99 into other cellular compartments. The dynamics of HA-C99 distribution in lysosomes were similar under different conditions. The two constructs carrying mutations in the intracellular domain, TM/IC(ala)KNKTG and TM/IC(ala)EWV, differed mainly from pcDNA (Fig. 3F, 4E), with an increase in HA-C99 colocalization with lysosomes of 18% (LogBF10 = 0.97) and 21% (LogBF10 = 5.27), respectively. Although TM/IC had low LogBF10 values compared to pcDNA (LogBF10 = 0.13), it increased HA-C99 in lysosomes by 18%, which is similar to TM/IC(ala)KNKTG and TM/IC(ala)EWV (18% and 21%, respectively), whose LogBF10 values are consistent. Thus, the presence of MT5-MMP TM and IC (intact or mutated) increases the presence of HA-C99 in lysosomes.

Overall, MT5-MMP variants that preserve high levels of C99 and secreted Aβ (i.e., TM/IC and TM/IC(ala)EWV), were more efficient at reducing HA levels in the Golgi while maintaining higher levels in the ER, compared to variants that reduce both C99 and Aβ (e.g., EXT and TM/IC(ala)KNKTG). Furthermore, MT5-MMP TM and IC appear to play an important role in the subcellular sorting of HA-C99, promoting its exit from the Golgi and its translocation into lysosomes.

### MT5-MMP modifications alter colocalization with HA-C99

Based on the previous data, MT5-MMP variants (Fig. 5A) likely alter the fate of C99 by affecting its distribution within the endomembrane system. However, whether these changes are driven by proximal interactions between MT5-MMP variants and C99 remains unclear. To address this, we performed a proximity ligation assay (PLA) using our RPE1 cell-based model^19^. PLA detects when two target proteins are within <40 nm of each other. Here, HA-C99 and MT5-MMP were targeted using antibodies against their N-terminal HA and Flag tags, respectively. For quantitative analysis, we used pcDNA as a reference control and subtracted the number of positive spots detected in pcDNA from those observed in the MT5-MMP variants, considering pcDNA spots as the baseline for nonspecific interaction.

**Fig 5.**
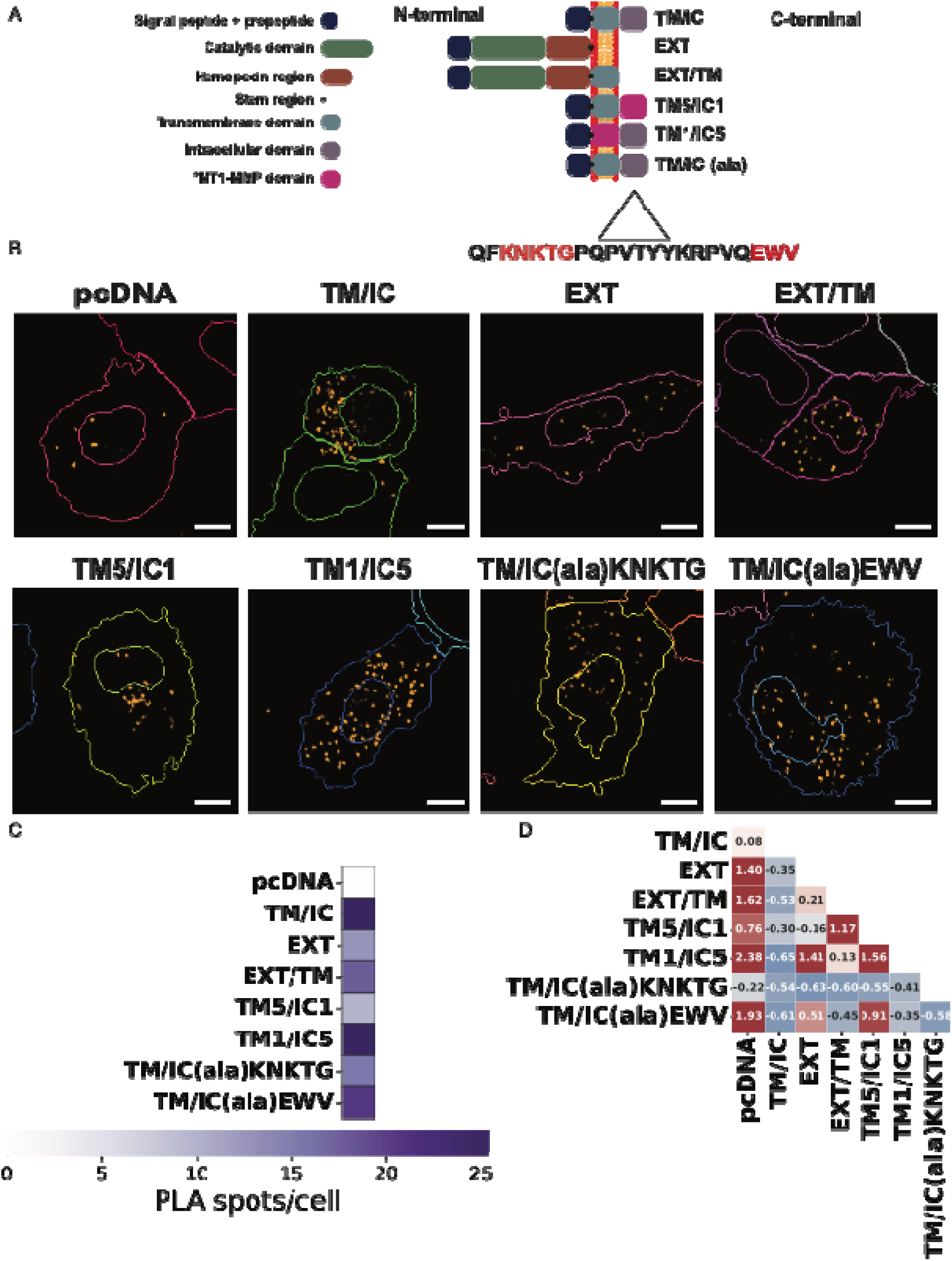
Mutating residues in the MT5-MMP IC domain disrupts interactions with C99. (A) Schematic representation of the MT5-MMP variants used. The amino acid groups in red (KNKTG and EWV) were replaced by ala. (B) Representative high content screening (HCS) microphotographs showing segmented (nuclear and cytoplasmic borders) RPE1 cells co-transfected with HA-C99 and the MT5-MMP variants described in (A). Orange dots correspond to proximity ligation assay (PLA) spots reflecting the proximity of HA (HA-C99) and Flag (MT5-MMP variant) at less than 40 nm, revealed by antibodies conjugated to DuoLink© orange oligos. The scale bar is 10 µm. (C) Quantification of the number of PLA^+^ spots in each MT5-MMP variant. PLA^+^ spots in pcDNA were subtracted from the other conditions, as we considered them to represent the non-specific background signal. Values representing a high number of PLA^+^ spots are purple and darker, while lower PLA^+^ spots are represented in lighter purple colors or white. (D) Diagonal correlation matrix representing the LogBF10 values for each comparison in a Bayesian ANOVA with post-hoc multiple comparisons of the number of PLA^+^ spots between different MT5-MMP variants. Higher values are represented in light-to-dark red, and lower values are represented in light-to-dark blue. For instance, the difference in PLA^+^ spots between TM1/IC5 and TM5/IC1 has a LogBF10 value of 1.56, suggesting that there is a difference in the average number of PLA+ spots per cell between these two variants.

Quantification of HA-C99 colocalization differed across all MT5-MMP variants compared to pcDNA. Individually, TM/IC and TM1/IC5 cells exhibited the highest number of PLA+ spots (25 spots each per cell) (Fig. 5B, C). Despite absolute differences, TM/IC showed low LogBF10 values in individual comparisons with other constructs (Fig. 5D). TM1/IC5 (25 spots) showed increased colocalization with HA-C99 compared to the TM5/IC1 chimera (10 spots, LogBF10 = 1.56) and EXT (13 spots, LogBF10 = 1.41). TM/IC(ala)EWV (21 spots) was the third construct with a high number of PLA^+^ spots, exceeding the 13 spots of EXT (LogBF10 = 0.51) and the 10 of TM5/ICI (LogBF10 = 0.91). Finally, compared to EXT/TM (18 spots), TM5/IC1 (10 spots) showed reduced colocalization with HA-C99 (LogBF10 = 1.17) (Fig. 5 A-C). Taken together, these results indicate that the MT5-MMP IC domain plays a key role in mediating the colocalization (and potentially the interaction) of the proteinase with HA-C99. However, KNKTG or EWV substitutions were insufficient to significantly decrease the proximal interactions of MT5-MMP with C99, suggesting that these IC mutants retain the ability to interact with C99 while at the same time driving its translocation to other subcellular compartments.

### MT5-MMP-based peptides promote C99 degradation

The ability of (ala)KNKTG to reduce C99 and Aβ levels, along with its capacity to colocalize with C99, suggests its potential as a framework for designing mimetic peptides that prevent C99 accumulation. To test this hypothesis, we synthesized MT5-MMP IC peptidomimetics containing mutations that reduced C99 levels when overexpressed with plasmid transfection. We generated two peptides: p17 (KNKTG substitution) and p20 (PVQ substitution). The peptides were co-transfected with HA-C99 in HEK cells for 48 h, with no cytotoxic effect detected by MTT assay (Fig. 6A, B). Treatment with p20 resulted in a significant 38% decrease of HA-C99 levels only at the highest concentration of 1000 nM (Fig. 6C). In contrast, p17 induced a dose-response reduction in HA-C99 levels at 1, 10, 100 and 1000 nM, reducing HA-C99 by 29%, 35%, 53%, and 41%, respectively, with statistically significant at the latter three concentrations (Fig. 6D). Given its higher efficacy, we developed a pilot assay to evaluate the potential of p17 conjugated to a vector enabling intracellular delivery without transfection. We conjugated p17 to C5 (C5p17), a VHH (nanobody) that binds to the transmembrane transferrin receptor (TfR), facilitating endocytosis and internalization^20^. We confirmed the proof of concept: transfection of the VHH-p17 conjugate maintained the reduction of HA-C99 levels by 37%, 37% and 36% in HEK cells at 1nM, 10 nM and 100 nM, respectively (Fig. 6E).

**Figure 6.**
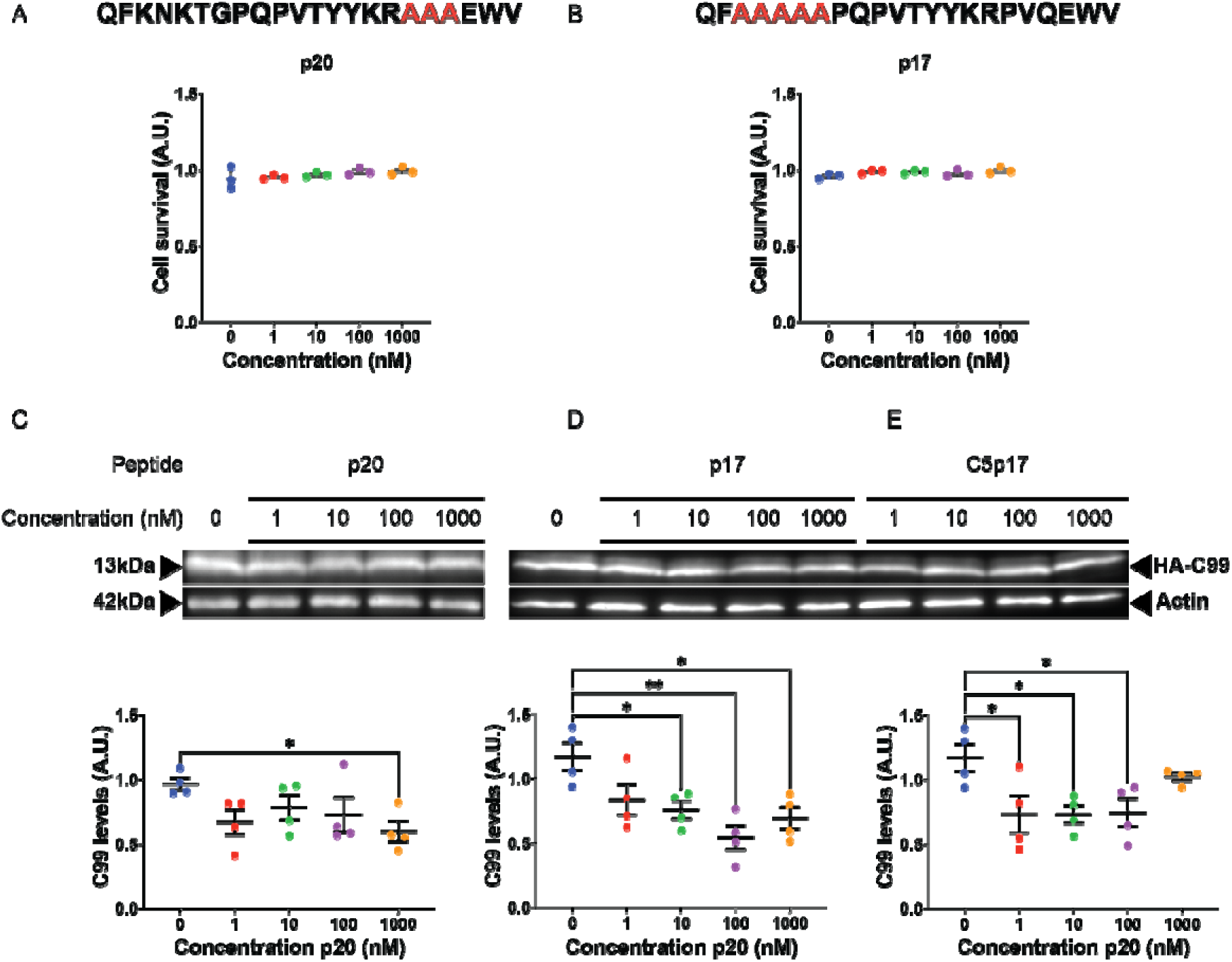
Effects of modified peptides of the MT5-MMP IC domain on C99 levels in heterologous cell lines. (A-B) Amino acid sequence of peptide p20 (A) and peptide p17 (B), based on the MT5-MMP IC, where PVQ and KNKTG were substituted by ala (in red). The graphs below show the absence of cytotoxicity measured by the MTT assay 48 h after cotransfection with C99 and p20 (A) or p17 (B). Absorbance values are expressed in arbitrary units (A.U.) and normalized to those of samples treated only with transfection reagent alone. (C-E) Representative western blots of HEK cells co-transfected with HA-C99 and p20 (C), p17 (D) or p17conjugated conjugated with a C5 VHH (C5p17) (E) at different concentrations. HA-C99 was detected with a rat anti-HA antibody. The graphs below show semi-quantifications of HA-C99 levels normalized to actin as a protein loading control. Values are expressed as mean ± SEM. P-values are represented by * p < 0.05 and ** p < 0.01. ANOVA followed by Dunnett’s post hoc multiple comparison test, with at least three independent experiments.

## DISCUSSION

In recent years, we have demonstrated the pathological role of MT5-MMP in cellular and animal models of AD^8,9,21^, as well as the critical importance of its non-catalytic TM and IC domains in regulating C99 and Aβ accumulation^4^. Building on these findings, the present study identifies key mutations in the TM and IC domains of MT5-MMP that modulate its functional interactions with C99, thereby altering its processing and intracellular distribution. Chimeric combinations of MT5-MMP and MT1-MMP TMs and IC domains revealed distinct proximity interactions and subcellular localization patterns that differentially influence C99 and Aβ levels. In addition, alanine substitutions of KNKTG or PVQ clusters in the IC domain reduced C99 levels, with only the KNKTG substitution consistently decreasing Aβ accumulation. Based on these results, we generated and vectorized a synthetic peptide mimicking the MT5-MMP IC KNKTG mutation, which effectively reduced C99 levels, reproducing the effect observed with plasmid transfection of this MT5-MMP variant. Overall, these findings identify specific amino acid clusters in the TM and IC domains as critical regulators of C99 fate and demonstrate the feasibility of leveraging these modifications to develop anti-C99 therapeutic agents.

### Modifications in MT5-MMP domains alter C99 fate

Previous studies have shown that the MT5-MMP TM/IC variant, when coexpressed with C99 in HEK cells, replicates the pro-amyloidogenic effects of full-length MT5-MMP, increasing both C99 and Aβ levels. In contrast, MT5-MMP variants lacking the TM/IC or IC domains significantly reduce in C99 and Aβ, highlighting the role of the C-terminal domains in regulating C99/Aβ metabolism^4^. The catalytic inactivation of MT5-MMP extracellular domain (ΔEXT) did not prevent the reduction in C99 levels—primarily mediated by proteasome activity—, but restored Aβ levels. This suggests that while MT5-MMP proteolytic activity is not involved in C99 degradation, it may influence Aβ metabolism, as previously reported for its homologue MT1-MMP^4,22,23^.

MT1-MMP and MT5-MMP share approximately 60% overall homology, yet their TM and IC domains exhibit only ∼20% sequence similarity^24^. This divergence may explain the opposing effects of MT5-MMP and MT1-MMP variants lacking the IC domain: MT5-MMP IC deletion decreases C99 levels, whereas MT1-MMP IC deletion increases them in HEK cells^4^. Consistent with these findings, swaping TM and IC domains between the two MT-MMPs differentially regulated C99 and Aβ levels. Thus, the accumulation of C99 and reduced Aβ levels in cells expressing TM5/IC1 suggest that it protects C99 from both proteasomal and γ-secretase processing, as neither pathway inhibitor (i.e., LACT and D6) restores C99 levels. In contrast, TM1/IC5 chimera facilitated C99 processing, as proteasome and γ-secretase inhibition rescued C99 accumulation. Together, these results indicate that specific TM/IC domain combinations may differentially regulate amyloidogenesis by modulating C99 stability and its accessibility to proteasomal and γ-secretase pathways.

### Specific IC domain mutations prevent C99 and Aβ accumulation

To probe the functional significance of sequence divergence between MT5-MMP and MT1-MMP, we introduced deletions or alanine substitutions in specific amino acid clusters of the MT5-MMP IC domain. While most deletions had negligible effects, KNKTG deletion significantly reduced C99 levels. This effect was replicated and even enhanced by alanine substitutions of KNKTG or PVQ, with only the KNKTG substitutions reducing also Aβ levels. This suggests a pivotal role for amino acids adjacent to the TM domain in regulating C99 and Aβ metabolism. The inability of LACT and D6 to restore C99 levels in cells expressing TM/IC(ala)KNKTG implies the activation of alternative, proteasome- and γ-secretase-independent degradation pathways for C99. This contrasts with previous reports where overexpressed C99 was primarily processed by the proteasome^4,25^, emphasizing the unique regulatory role of the TM/IC(Ala)KNKTG substitution in regulating.

In contrast, alanine substitutions in the C-terminal PDZ-binding EWV motif not only maintained C99 and Aβ levels comparable to those in TM/IC controls but also enhanced γ-secretase-dependent C99 processing, as demonstrated by the increase in C99 levels upon D6 treatment. Meanwhile, both LACT and D6 rescued C99 levels in the TM/IC control cells, suggesting that EWV substitution disrupts MT5-MMP interaction Mint3, a protein known to regulate MT5-MMP subcellular trafficking^26^. Given that Mint3 also binds to the APP C-terminal domain and modulates its intracellular traffic^27,28^, the 45% reduction in C99 localization in the Golgi observed with EWV substitution may reflect disrupted MT5-MMP subcellular localization. This could facilitate the export of C99 to acidic subcellular compartments where γ-secretase is more active^29–31^, thereby promoting its conversion to Aβ.

Collectively, these results demonstrate that subtle modifications in the IC domain of MT5-MMP redirect C99 into distinct proteolytic pathways, likely by altering its distribution. This underscores the remarkable plasticity of the MT5-MMP IC sequence in modulating C99 metabolism and reinforces its potential as a framework for developing anti-C99/Aβ strategies.

### MT5-MMP variants alter C99 subcellular localization

Our findings suggest that MT5-MMP variants induce changes in the subcellular distribution of C99, potentially reflecting shifts in its processing pathways. This is consistent with the role of canonical secretases in regulating the subcellular localization of C99/APP and Aβ production^32–34^.

Principal component analysis (PCA) revealed differential effects of MT5-MMP variants on HA-C99 distribution, highlighting the importance of the TM domain and the C-terminal EWV motif. Cluster analysis of the sum of LogBF10 values confirmed that TM/IC(Ala)EWV most strongly influences HA-C99 distribution across the endomembrane system, whereas other MT5-MMP variants exert more localized effects. This underlines the importance of the EWV motif in regulating the subcellular localization of C99, likely by preventing its interaction with Mint proteins and facilitating γ-secretase cleavage. Beyond Mint3, other Mint family members interact with APP metabolites, and Mint1 overexpression or its PTB domain has been shown to decrease Aβ40 secretion by inhibiting γ-secretase cleavage of C99 in HEK cells^35^. Thus, EWV mutations may enable C99 to bypass Mint1-dependent inhibition of γ-secretase cleavage.

Individual analysis of each variant revealed a general trend toward reduced HA-C99 levels across organelles, particularly in the endoplasmic reticulum (ER), the main site of protein synthesis. This is consistent with the stimulation of C99 sorting to other compartments. With respect to the Golgi apparatus, TM/IC(Ala)EWV induced the greatest reduction in C99 levels (−45%), likely due the disruption of MT5-MMP recycling to the plasma membrane^26^, thereby enhancing TM/IC(Ala)EWV influence on C99 intracellular sorting.

Expression of the TM/IC(Ala)KNKTG and TM/IC(Ala)EWV variants significantly increase C99 in lysosomes. Given their opposing effects on C99 and Aβ levels—KNKTG acting as a depressant and EWV as a stabilizer—this finding suggests that alterations in C99 intracellular trafficking prior to lysosome translocation. Recent studies have evidenced intracellular lysosome heterogeneity, supporting the idea that C99 may traffic to molecularly distinct lysosomes^36^. In this line, the inhibition of the autophagosome pathway increases C99 levels in LAMP1+ lysosomes in human neuroglioma cells, whereas autophagosome activation has the opposite effect and prevents γ-secretase cleavage of C99^37^. Late endosomes and lysosomes are also major sites of C99 conversion into Aβ^31^. KNKTG and EWV mutations in MT5-MMP IC domain distribute C99 into distinct cellular compartments, thereby directing it to heterogenous lysosomes containing different proteolytic pathways. In addition, TM/IC(Ala)EWV-mediated increase in lysosomal C99 could initiate a pathogenic loop wherein elevated C99 elevated disrupt Ca^2+^ homeostasis in late endosomes and lysosomes, further promoting C99 accumulation^38^.

In summary, different MT5-MMP variants distinctly affect C99 subcellular distribution. Chimeric combinations or minimal mutations in the MT5-MMP IC domain redistribute C99 within the endomembrane system, underscoring the key role of the C-terminal domains in C99 trafficking.

### IC domain mutations determine MT5-MMP colocalization with HA-C99

Given that MT5-MMP and C99 coimmunoprecipitate when overexpressed in HEK cells^4^, changes in C99 subcellular distribution may result from altered proximal interactions with MT5-MMP variants. High-throughput PLA screening revealed that TM/IC and TM1/IC5—both containing intact MT5-MMP IC—exhibited the highest level of interaction with HA-C99, in contrast with the low PLA signal observed in TM5/IC1-expressing cells. Together, these data suggests that the IC of MT5-MMP facilitates colocalization and interaction with C99, consistent with the sufficiency of MT5-MMP TM/IC to coimmunoprecipitate C99 in HEK cells^4^. The strong reduction of PLA signals upon expression of EXT further supports the importance of the MT5-MMP IC domain for proximal interaction with C99. However, the increased colocalization of EXT/TM with C99 compared to EXT alone indicates that MT5-MMP TM also contributes to this interaction. Together with the decreased PLA signal in TM5/IC1, these data indicate that the interface between the TM and IC domain is important, though not exclusive, for maintaining proximal interactions between MT5-MMP and C99. Further structural analysis of the MT5-MMP TM and IC domains could provide additional insights on the mechanisms underlying these dynamics.

Notably, the KNKTG or EWV substitutions in the MT5-MMP IC maintained colocalization with C99, indicating that these modifications alter C99 processing without disrupting TM/IC interactions with C99. Thus, TM/IC(Ala)KNKTG could serve as a reference sequence for designing molecular tools that interact with C99 to modulate its the subcellular trafficking and degradation, ultimately decreasing C99 and Aβ levels.

### A synthetic MT5-MMP IC peptide mimicking s KNKTG mutation reduces C99 accumulation

Synthetic peptides are widely used as relevant therapeutic tools^39^, and offer a promising platform for exploiting the anti-C99 properties of modified MT5-MMP IC domains. In this study, the synthetic peptide containing the (Ala)KNKTG sequence significantly reduced C99 levels, while the peptide carrying the (Ala)PVQ substitution (p20) was less effective. These data provide initial proof of concept for the feasibility of MT5-MMP IC-based strategies to promote C99 degradation, likely through mechanisms that alter C99 cellular trafficking. In this vein, a recent study showed that editing the YENPTY motif C99— a sequence critical for its trafficking—resulted in a sustained reduction of C99 and Aβ levels in an AD knock-in mouse model^40^. This highlights the importance of targeting APP metabolites with strategies that influence their trafficking and eventually the production of toxic biomolecules. Our study adds to this concept by demonstrating the potential of using MT5-MMP as a benchmark to generate synthetic peptides, a generally safe therapeutic strategy that also avoids genome modification, while effectively promoting C99 and Aβ clearance in the context of AD.

### Conclusion

In this study, we highlight the importance of specific amino acid clusters in the non-catalytic TM and IC domains of MT5-MMP to regulate C99 and Aβ metabolism. Sequence modifications in these domains alter MT5-MMP colocalization with C99 and influence the subcellular trafficking and processing of C99, with distinct impacts on C99 and Aβ levels. These findings provide new mechanistic insights into the complex pathways governing APP metabolite homeostasis and highlight the potential of targeting non-catalytic functions of MT5-MMP. Moreover, we show that engineered MT5-MMP variants and IC-derived peptidomimetics may serve as original tools to promote C99 and Aβ clearance, offering novel avenues for therapeutic intervention in AD.

## Declarations

### Ethics approval

All the experimental procedures were conducted in agreement with the authorization by the French Ministry of Research to the laboratory (# 7950), as defined in the directive 2009/41/CE of the European Parliament and the European Council of May 6 2009 for the use of genetically modified organisms.

### Availability of data and materials

All the data generated or analyzed during this study are included in this published article and are available from the corresponding author upon reasonable request. All the analysis files in the form of Jupyter Notebook are uploaded and freely accessible at: github.com/RiveraLabINP/BelioMairal_2024

### Credit authorship contribution statement

**Pedro Belio-Mairal**: Conceptualization, Methodology, Software, Validation, Formal analysis, Investigation, Data curation, Writing – original draft, Writing – review & editing, Visualization. Funding acquisition. **Athina Kamitsou**: Methodology, Formal analysis, Investigation. **Delphine Stephan**: Methodology, Investigation. **Nicolas Jullien:** Methodology, Investigation, Resources. **Diarra Thiane:** Investigation. **Laurence Louis**: Investigation. **Florian Benoist:** Investigation, Resources. **Baptiste Serrano**: Resources. **Marion David:** Resources. **Michel Khrestchatisky**: Resources, Writing – review & editing. **Pascaline Lécorché:** Resources, Writing – review & editing. **Emmanuel Nivet:** Conceptualization, Supervision, Writing – review & editing, Project administration. **Santiago Rivera**: Conceptualization, Methodology, Validation, Investigation, Data curation, Writing – original draft, Writing – review & editing, Visualization, Supervision, Project administration, Funding acquisition.

## Acknowledgements

We thank F. Chechler and S. Gon for generously providing HEK cells and RPE-1 cells, respectively. We thank H. Kovacic and F. Parat for their technical advice, and R. Pardossi-Piquard, N. Sergeant, and S. Claeysen for their insightful suggestions on the manuscript.

